# Focality of sound source placement by higher (9^th^) order ambisonics and perceptual effects of spectral reproduction errors

**DOI:** 10.1101/2024.08.07.606870

**Authors:** Nima Zargarnezhad, Bruno Mesquita, Ewan A Macpherson, Ingrid Johnsrude

## Abstract

Higher-order ambisonic rendering is an increasingly common soundscape reproduction technique that, in theory, enables presentation of virtual sounds at nearly any location in 3D space, relatively unconstrained by the veridical locations of loudspeakers. We evaluated whether 3D sound reproduction through a 9^th^-order ambisonic loudspeaker array was indeed sufficiently accurate to probe the limits of human spatial perception. We first estimated minimum audible angles for human listeners for a variety of reference points on the horizontal plane. We demonstrated that the system can reproduce sounds with a spatial resolution that is equal or superior to the limits of human acuity, at least on the horizontal plane at the front. Importantly, the resolution of ambisonic reproduction appeared equivalent for regions of the system with high and low loudspeaker density. We also estimated localization cues at the same locations and showed that although localization cues for low-frequency components were well preserved, they were somewhat distorted for components above 4000 Hz. Finally, we provide evidence that these high-frequency distortions can serve elevation cues by human listeners. In summary, we showed that a 9^th^ order ambisonic system is able to render highly focal sound sources, and that high-frequency reproduction distortions may introduce unwanted localization cues.

## I. INTRODUCTION

### A. Background

Human auditory perceptual experience is essentially spatial in nature, despite the fact that, unlike for vision and touch, the sensory epithelium is not spatiotopically organized. Our skin and retinas are equipped with receptors that, by their location in the sensory array, encode information about the spatial position of events in the world. Our cochlea, however, is organized by frequency, and appears to compute location from a variety of different temporal and spectral sound features often requiring integration across the two ears. Sound location relative to the head is primarily estimated through binaural cues, such as Interaural Time Difference (ITD) and Interaural Level Difference (ILD) which index the delay of sound arrival time and level between ears^1,2^. While spectral detail in reasonably broadband sounds provides cues to elevation, cues for horizontal localization are provided by ITDs (for lower frequencies; below 1000 Hz) and ILDs (more salient at higher frequencies; above 1500 Hz)^1,3,4^. In addition to these cues, which inform the listener about the lateral location of a sound source relative to the midline, monoaural spectral cues contribute to estimating the elevation and front/back location of the source. These cues are the result of sound filtering by the listener’s head, upper body, and by the pinnae of the ears. The Head-Related Transfer Function (HRTF), is an individual response function that characterizes the sound received by each ear. A pair of HRTFs for a given individual can be used to filter a sound such that it is perceived by that person as coming from a particular location in 3-dimensional (3D) space. Individual HRTFs enable the recreation of a spatial hearing experience in experimentally controlled settings, over headphones^1,5–7^. In practice, because estimating individual HRTFs is a time-consuming, expensive, complex (requiring specialized equipment) and somewhat unpleasant process (usually involving the insertion of a microphone near the eardrum), generic HRTFs are typically used^2,7^. These simulations, while delivering a ‘spatialized’ percept, generally do not deliver as compelling a spatial percept as free-field listening.

High-density 3D LoudSPeaker (LSP) arrays are expensive, but deliver a compelling, free-field, spatial percept, capitalize fully on an individual participant’s own HRTFs, and afford opportunities to study naturalistic behaviors (such as audio-visually guided grasping). In the basic implementation, each sound source is presented from exactly one LSP. This method, Single-Channel (SC) presentation, limits the presented sound source locations to the locations of actual LSPs - which are typically many centimeters apart. This is obviously inadequate for the study of the limits of human spatial perception. Another method, called Vector Based Amplitude Panning (VBAP), overcomes the limited resolution of SC presentation by distributing a virtual sound source’s energy among the surrounding LSPs and enables the simulation of sound sources between LSP locations. VBAP first determines the minimum number of nearby LSPs for any given location then distributes the sound gain among those LSPs such that their vector sum results in a sound image with the appropriate gain at the intended location^8^. This algorithm essentially utilizes one LSP when the intended sound source coincides with an LSP location (identical to SC presentation in this situation); two LSPs when it aligns on the line between two LSPs; and three LSPs if the sound location neither coincides with an LSP nor aligns with the line connecting two LSPs. The energy spread of a sound source using VBAP presentation depends on its distance from nearby LSPs and LSP geometry, and the number of LSPs utilized to reproduce it. The energy spread of sound sources determines their perceived spatial occupancy; for example, sound sources with larger energy spreads are perceived to occupy larger portions of the space (i.e., they are blurred in space). Therefore, sound source images are nonhomogeneous in their apparent diffusion^9^.

Ambisonic panning is another reproduction method that utilizes a 3D LSP array to reproduce 3D soundscapes. For ambisonic panning, the soundscape (comprising one sound, or many) is decomposed into a series of spherical harmonics that describe the sound pressure level, velocity, and its higher order derivatives in different directions within the 3D acoustic environment. Although utilizing higher order derivatives of the sound wave velocity enables the encoding of more focal sound sources, decoding them in the virtual acoustic environment requires more presentation channels (LSPs); therefore, the spatial resolution of ambisonic reproduction is limited by the number of LSPs in the array. Theoretically, ambisonic panning has higher spatial resolution than SC and VBAP presentation methods with an equalized energy spread using all LSPs for all sound source positions^10^. With lower order ambisonic panning systems (orders 1 to 4), rendered sound sources are perceived as “blurry” or diffused, and resolution is poor. As the order of the system increases, rendered sound sources are, in theory, less blurry, such that a normal listener might perceive them as more crisply focal and thus be better able to resolve closely adjacent sources^11^.

Generally, all sound reproduction methods are an approximation of real-world acoustic characteristics and reproduction is not perfect. The perceptual effects of reproduction errors and the extent to which these errors are acceptable depend on the intended use of the system. Presenting sound sources with SC will minimize errors in localization, but it limits the spatial resolution of soundscape reproduction. VBAP has greater spatial resolution than SC, but the energy of sound sources is spread unequally throughout the 3D space, such that the blurriness of sound sources depends on their location relative to LSPs. Although rendering through ambisonics improves on SC and VBAP, even higher order systems may not perfectly reproduce sound locations: errors in timbral and reconstructed ITD and ILD information have been noted in simulations^12^. Additionally, soundscape reconstruction with ambisonics is valid within the area in which the superposition of spherical harmonics is reliable; a minimum ambisonics order is required for the system to reproduce planewaves below a particular wavelength reliably, within a sphere of a particular size located at the center of the array^6,13^. The possibility of reproduction errors affecting perception may preclude the use of this technology for spatially precise reproduction of acoustic environments.

The goal of the present study is to validate the utility of a custom-built LSP array for conducting studies of human perception of auditory space. We aim to explore how accurately and precisely human listeners can localize sound sources with 9^th^ order ambisonic panning to examine whether the spatial resolution of this system – its ability to reproduce highly focal sound sources in specific locations – is sufficient to challenge the limits of human spatial acuity. Neither single-channel presentation nor VBAP can accomplish this easily for an extended region of space. Furthermore, we hope to identify the limitations of our particular ambisonic system for those who conduct experiments with this technology, although the work is also valuable for others using similar Higher-Order Ambisonic (HOA) arrays for human auditory perception research. Future system designers may also benefit from our observations.

### B. The AudioDome

All the experiments in this manuscript used a custom-made 3D LSP array at Western University, called the “AudioDome” (FIG 1.A, sonible, GmbH, Austria) for sound presentation. The AudioDome consists of 4 dual-channel subwoofers and 91 LSPs arranged in a geodesic-like sphere with a radius of approximately 1.65 m, located in a hemi-anechoic room (FIG 1.A). The system was built with KEF E310 LSPs with 4.25” coaxial drivers in oval enclosures, with approximately linear frequency response (90 Hz-33 kHz, ± 3dB). The subwoofers have 2 independent drivers and were installed to assist with the lower portion of the spectrum (<150 Hz). Although all the audio files of our experiments had some content below 150 Hz we did not use the subwoofers as we were interested in focal sound source reproduction at a higher elevation. The overall bandwidth of the Sonible system was estimated by playing a linear sine sweep from 0 Hz to 22050 Hz. This was recorded with an Earthworks M30 measurement microphone (30-kHz bandwidth) at 48000 Hz sampling rate and demonstrated that the frequency range used in our acoustical analyses (100-20,000 Hz) was within the 3-dB bandwidth of the system.

**FIG. 1.**
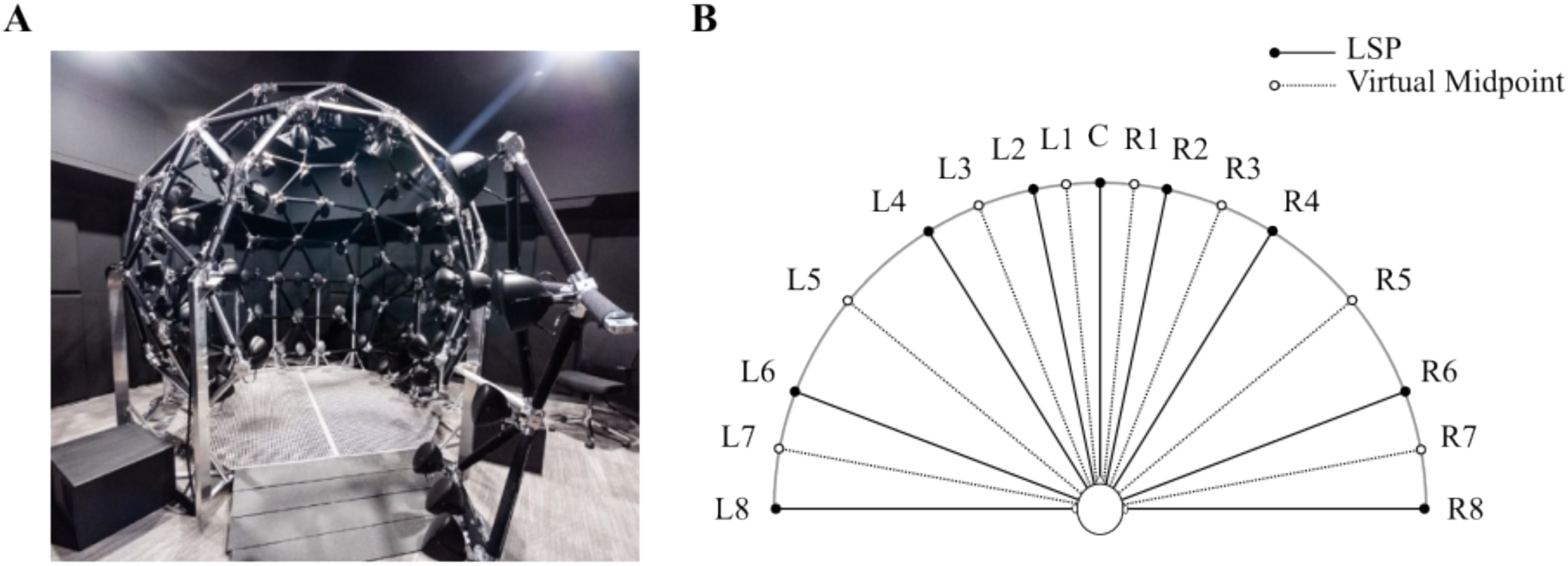
(color online) A. The AudioDome (sonible, GmbH, Austria): the array consists of 91 LSPs and 4 subwoofers arranged in a spherical shape with a radius of approximately 1.65 m in a sound attenuating chamber. This system is equipped with SC presentation, VBAP, and 9^th^ order ambisonic panning technologies. (The AudioDome doorway is open in this picture and therefore some LSPs are not in position.) B. A schematic of the reference points used in Experiments 1-3. Locations of LSPs (filled circles) and locations between LSPs (open circles) on the horizontal plane with their labels and the participant’s head at the center.

The chamber dimensions are 5.75 m×5.75 m×3.40 m, and the AudioDome’s center is located 2.34 m away from the side walls. All the surfaces on the chamber walls and ceiling are covered with black high-density glass wool panels encapsulated with micromesh and fabric coating mounted on a custom-made self-supported aluminum rig and high-performance diaphragmatic resonators installed on the ceiling and the walls. The floor is carpeted and there is some acoustic foam under the AudioDome’s grid floor which is approximately 12 cm above the carpet. Black Velcro was used to cover the frame supporting the LSPs in order to minimize potential acoustic and optical reflections. (Optical reflections provide visual information that might confound sound localization experiments^14^. The material is black to absorb any unintended light energy.) The room’s impulse response was measured via the exponential sweep method using the iPad-based “Audio Tools” application (Studio Six Digital). The excitation loudspeaker was placed near an AudioDome loudspeaker, and the measurement microphone was at the central listening position. Analysis of the impulse response revealed that 1) the wideband reverberation time (60-dB decay time estimated via RT30) was 96 ms, 2) the largest octave-band RT30 was 127 ms in the 125-Hz band, and 3) some “clutter” reflections from the AudioDome speakers were evident in the impulse response from 5-11 ms following the direct sound, but with peak levels at least 18 dB below the direct sound.

The AudioDome is capable of SC presentation, VBAP, and 9^th^-order ambisonic panning to reproduce 3D acoustic environments. For all presentation/panning methods, .wav or .mp3 file formats of single auditory objects can be fed into AudioDome’s Spatial Audio Creator software input to be decoded and encoded appropriately (in case of ambisonic presentation, always encoded and decoded in 9^th^ order). Additionally, sound-field recordings with microphone arrays can be sent as ‘atmosphere’ files to the software input to be projected and presented appropriately. In the experiments presented in this manuscript, we only used .wav audio files generated and presented at 44100 Hz sampling rate.

### C. Overview of the Experiments

We first measured horizontal spatial acuity by human listeners, in order to evaluate whether 9^th^ order ambisonic panning can reproduce sounds with sufficiently focused spatial resolution that they can be resolved at the very limit of human spatial acuity in the horizontal plane. Traditional SC presentation has determined that Minimal Audible Angles (MAA) are ∼1° on the horizontal midline, and then acuity monotonically decreases to ∼2° at 45° azimuth, and to more than ∼8° at azimuth locations at or above 75° off the midline.^1,15,16^ In Experiment 1, we measured listeners’ horizontal MAAs. We also investigated whether obtained MAAs depended on the distance of the rendered source from the nearest LSPs: higher MAAs in virtual positions further away from LSPs would indicate that the focality of rendered sources was not the same everywhere on the surface of the sphere defined by the LSP array.

Subsequently, using a Head-and-Torso Simulator (HATS), we compared the spectral composition of rendered sound, and reconstructed ITD and ILD cues delivered to an example listener at the center of the array (Experiment 2), for SC presentation, VBAP and ambisonic panning.

Finally, based on the findings in the first two experiments, we measured the effects of spectral distortions introduced by ambisonics on the perception of elevation by human listeners (Experiment 3).

## II. EXPERIMENT 1

The first experiment examined whether the spatial blurring of sound sources reproduced with 9^th^-order ambisonics was above or below the threshold of human acuity (minimum audible angle) studied with SC reproduction^1,15,17^. Furthermore, we examined whether spatial acuity was worse (indicative of more diffuse, less focal sources) when virtual sources were located at a greater distance from the nearest LSPs.

We estimated Minimum Audible Angles (MAAs)^1,15,17^ on the frontal half of the horizontal plane at several locations that coincided with actual LSPs or at the midpoints of adjacent pairs (FIG 1.B). The highest spatial acuity (lowest MAAs) was expected at the front of the array (0° azimuth)^1,15^ and should approach 1° if ambisonic panning renders point sources at a spatial resolution at or below the limits of human acuity. However, the LSP density is also greatest at the location defined as 0° azimuth on the array (the location the participant habitually faces). Accordingly, we tested another condition in which participants were rotated to face another region of the array with the lowest LSP density, also on the horizontal plane. This allowed us to determine whether distance from the nearest LSPs changes the focality of sources. If we observe MAAs of ∼ 1° in both conditions, we can conclude that the array is able to render point sources at a spatial resolution at or below the limits of human acuity, everywhere on the surface of the sphere defined by the geodesic array.

### A. Materials and Methods

Participants were seven (4 female) young (aged 18-25 years), healthy, normally hearing (thresholds measured audiometrically at both ears at octaves of 125 Hz up to 8 kHz; participants were excluded if their threshold exceeded 25 dB HL at any tested frequency in either ear), right-handed, naive listeners with no history of neurological disorders. One participant who marginally passed the audiometry test was not able to achieve a high accuracy after three tries in the first session’s practice block and was not tested further. We collected and analyzed data from the remaining six participants. Participants sat in the middle of the AudioDome with their head positioned at the center of the dome. The chair was set to face the origin of the array’s coordinate system (0° azimuth and elevation angles) so that the ears aligned with the ±90° azimuth line, and the ear level was aligned with the 0° elevation level by adjusting the chair height.

Horizontal MAAs were estimated at 17 locations (“reference points”) on the frontal half of the horizontal plane. Nine of these coincided with LSPs within the ±90° azimuth range and eight were at midpoints between adjacent LSPs (FIG 1.B and TABLE I). Acuity (MAA) at each reference point was measured using a method of constant stimuli procedure, with 12 test points (6 to the left, 6 to the right) symmetrically located relative to each reference point, and a 13^th^ test point collocated with the reference point (see FIG 2). Each test point was paired with the reference point on 20 trials, such that the first noise burst of a trial was located at the “test point” on 10 trials, and was located at the “reference point” on the other 10 (i.e., the standard was presented first half the time). All 260 trials for a given reference point were presented successively, in a single block, with trials for different test points pseudorandomly intermixed such that the same test point was never presented on two successive trials. Thus, each participant was tested with 17 blocks of 260 trials each.

**FIG. 2.**
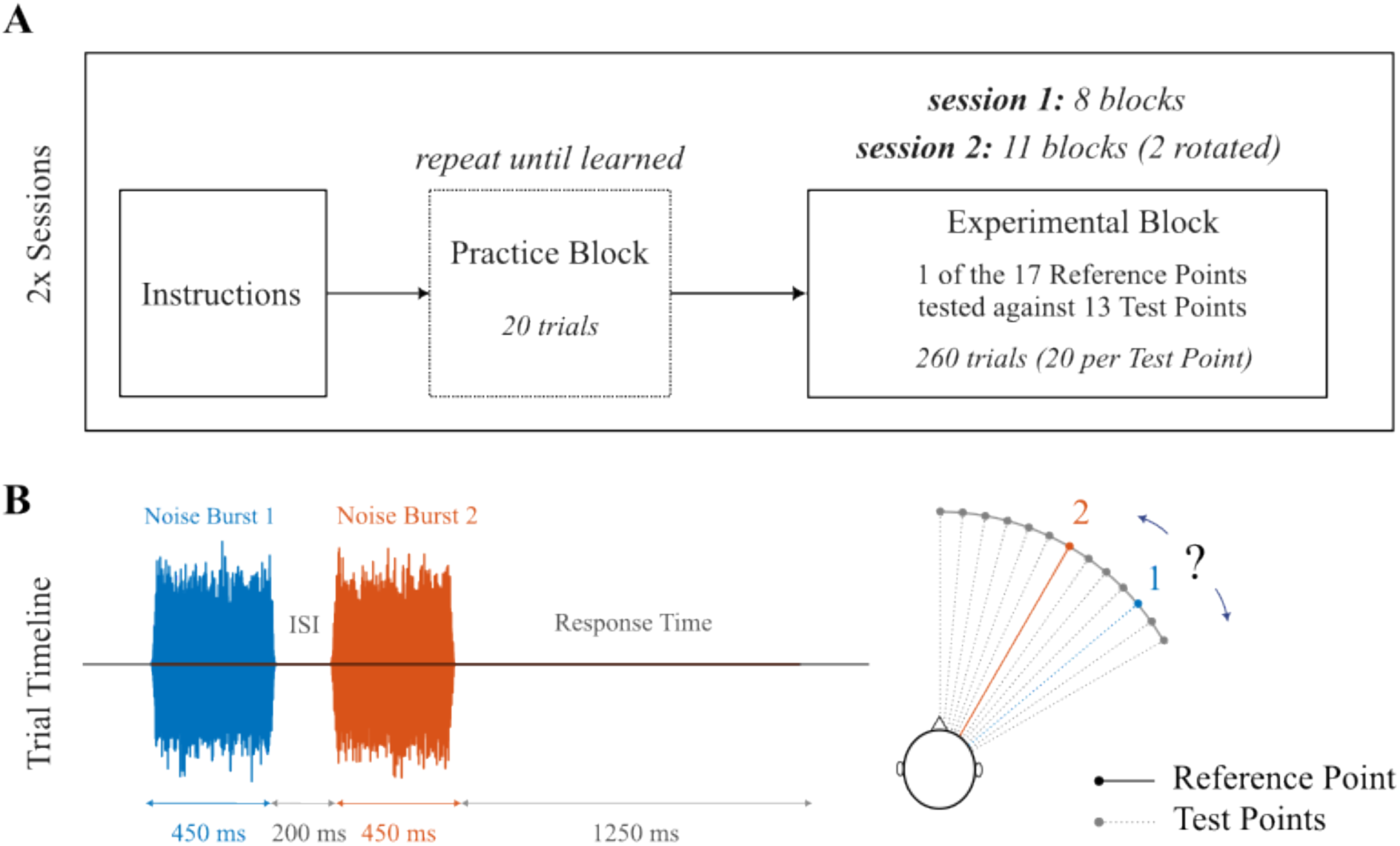
A. Experimental procedure: In the beginning of each of the 2 sessions participants were given the task instructions, which was to report whether the second sound of a pair was located clockwise, or counterclockwise, relative to the first. Participants then practiced the task over 20 trials in which single-channel presentation was used in the left horizontal plane of the AudioDome. Given the LSP spacing, distances were suprathreshold. The practice block was repeated until participants achieved perfect performance for such large location differences. Then they proceeded to the actual experimental blocks. In each block, one of the reference points was randomly chosen and tested against all 13 associated test points, 20 times each. Symmetrical reference points were tested within the same session. Both reference point C and the two ‘rotated’ reference points, when the participant’s chair was rotated such that L5, and R5, were on the midline, were tested in the second session. B. In each trial, a noise burst from the reference point and another burst from one of the test points were presented in a sequence. The bursts lasted 450 ms in duration, with a 200 ms gap between the pair. Participants had 1250 ms to report the relative direction of the second noise burst to the first. This duration was shown in pilot testing to be adequate for performance of the task (and was also used in the practice, described in the text, in which perfect performance was the criterion to advance to the main experiment). The reference point could be presented first or second. The right panel shows a schematic of a sample reference point, all its test points and the chosen test point for the trial illustrated in the left panel.

**TABLE I.** Reference point azimuth coordinate (φ), estimated average MAAs with standard errors (SE), and the number of participants remained in MAA estimation after excluding flat psychometric curves (N). Approximation of previously measured MAAs with pure tones in free-field are read from Fig. 6 of the Mills manuscript^15^ and put in the ‘Mills’ MAA’ column for comparison.

| Reference Point | | $\phi_{RP}$ (°) | | N | | MAA ± SE (°) | | Mills’ MAA (°) |
| --- | --- | --- | --- | --- | --- | --- | --- | --- |
| C | | 0 | | 6 | | $1.03 \pm 0.12$ | | ~1 |
| L1 | R1 | 5.82 | -5.82 | 6 | 6 | $1.11 \pm 0.08$ | $0.95 \pm 0.12$ | ~1 |
| L2 | R2 | 11.64 | -11.64 | 6 | 6 | $1.40 \pm 0.18$ | $1.29 \pm 0.17$ | ~1.2 |
| L3 | R3 | 21.68 | -21.68 | 6 | 6 | $1.64 \pm 0.12$ | $1.69 \pm 0.36$ | ~1.5 |
| L4 | R4 | 31.72 | -31.72 | 5 | 6 | $2.38 \pm 0.23$ | $2.10 \pm 0.32$ | ~1.8 |
| L5 | R5 | 50.41 | -50.41 | 6 | 6 | $4.46 \pm 0.66$ | $2.81 \pm 0.47$ | ~2 |
| L6 | R6 | 69.09 | -69.09 | 4 | 5 | $4.46 \pm 1.16$ | $8.48 \pm 1.55$ | ~3 |
| L7 | R7 | 79.55 | -79.55 | 4 | 5 | $5.84 \pm 2.30$ | $7.94 \pm 2.31$ | ~4 |
| L8 | R8 | 90.00 | -90.00 | 2 | 3 | $21.78 \pm 14.47$ | $6.24 \pm 3.46$ | > 10 |

On each trial, participants discriminated the location of the two stimuli (reference and test) by reporting whether the second noise burst of a pair was in a location clockwise, or counterclockwise, relative to the first burst. The probability of perceiving a test burst as clockwise from the reference burst was then plotted as a function of the distance of that burst from the reference point. The distance between test points was smallest near the midline, and larger off to the sides (i.e., the four most lateral reference points) given that acuity is poorer in this region^15,16^; test points ranged across ±12° from the reference for L8 and R8, ±9° for L7 and R7, and ±6° for all other reference points (to maintain the number of test points at 13, the increment size was adjusted accordingly, thus it was 2°, 1.5°, and 1° respectively). The MAA at each reference point was estimated by fitting a sigmoid to each individual’s psychometric data, and then reading off the 75% threshold value, which was taken as the just-noticeable difference (JND^18^). Equation 1 describes the model ψ_RP_(φ) used to describe the probability, ψ, that the test sound rendered at probe angle φ, was perceived to be counterclockwise relative to the sound rendered at the reference point, RP. Parameters α_RP_ and Δ_RP_, representing the center and steepness of the sigmoid function, were estimated using a non-linear least-squares method^19,20^.

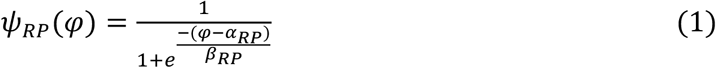

The two sounds (reference and test) were broad-band white-noise bursts (range 0-22050 Hz at 44100 Hz sampling rate presented via the 90-20730 Hz bandwidth system), 450 ms in duration, separated by a 200 ms silent gap^17^. Participants could respond (and change their response) up to 1250 ms after the second burst ended, as shown in FIG 2.B. The noise bursts were generated using the ‘wgn’ function in MATLAB 2022b software^21^ with the signal power set to −25 dBW with 22.5 ms long cosine^2^-shaped attack and decay ramps. The audio files were presented at 73.1 dBA to the participants in the AudioDome, which had a background noise level of 25.2 dB LAeq at its quietest (SNR = 47.9 dB). To avoid any effects of familiarity with particular tokens of noise on performance^22^ eighty different noise-burst tokens were randomly generated and pseudo-randomly assigned to different trials and intervals in the experiment (Each trial’s noise bursts were different). All eighty noise bursts had a flat spectrum and matched in duration and envelope.

The experiment was held in darkness to rule out the influence of vision on sound localization^14^; the only light source was the red LED used for eye fixation, placed at 0° azimuth and elevation. The LED was illuminated for the duration of each trial to inform the participant of the trial progression (illuminated at the start of each trial and extinguished at the end). The participants were instructed not to move, to keep their eyes open, and to fixate on the red light. An 8-mm magnetic sensor was attached to the back of the participants’ heads to track their motion. (Those head tracking data were not used). Experiment instructions did not make reference to the distinction between reference and test points. Instead, participants were instructed to make judgements on the position of the second sound they heard in relation to the first. On each trial, participants used two buttons on a small wireless keypad to indicate the direction in which they perceived the sound to ‘jump’ from the first burst to the second (clockwise/counter clockwise).

Reference point C, at 0° azimuth and elevation and which coincided with an LSP, is also surrounded by several other LSPs; four within ±11.64° azimuth and ±20.91° elevation. In contrast, reference points L5 and R5 are themselves virtual and in rather sparse LSP regions: the nearest LSPs are located at ±18.69° azimuth and±11.64° elevation (FIG 4.A). The higher concentration of LSPs at the front (reference point C) may result in more focal sound source reproduction at this location, enabling more accurate discrimination and lower MAAs compared to lateral positions, where LSPs are sparser. Human acuity is also best near the midline, and deteriorates laterally^1,15,16^. To ensure that LSP density is not affecting the focality of sound source reproduction (and hence acuity) we tested two further reference points (blocks): we rotated the chair so that participants faced L5 and R5. Now, these previously lateral positions, which do not coincide with an LSP and in which LSP density is low, are on the midline, just as reference point C was. This enables us to compare midline frontal MAAs for the reference points in the regions of highest (C) and lowest (L5 and R5) LSP concentrations. In these ‘rotated’ blocks, the participants completed the task for L5, and R5, as midline reference points. Thus, L5 and R5 were each tested twice as reference points; once as a more peripheral location, and once as a midline location. The red LED fixation point was set at L5 and R5 for these blocks.

The 19 experiment blocks (17 facing reference point C, one facing L5, and one facing R5) were split into two approximately 2 hours-long sessions over two days. Session One consisted of consent, audiometry, a demographic questionnaire, head measurements (not presented or analyzed here), followed by training, a practice block, and eight blocks testing four pairs of symmetric reference points. In Session Two, participants completed another practice block, then the remaining four pairs of reference points, the two rotated conditions and reference point C. At the end, the participants were asked about their attentiveness during the task, their understanding of the task, and any strategies they used to respond, then they were debriefed. The reference point pairs were randomly assigned to the two sessions at the beginning of the first session. The order of reference point blocks within a session was randomized. Also, the order in which the three frontal MAA blocks at C, L5, and R5 were tested was pseudo-randomly assigned to participants (each of the six possible permutations was assigned to a participant). Within each block, the order of the trials was pseudorandomized such that no test points were tested successively.

At the beginning of each session, participants completed a practice block which consisted of 20 trials with pairs of noise bursts presented at different locations drawn from the following locations: C, L2, L4, L6, and L8 (the left LSP locations). The leftmost noise burst of the pair was presented first on half of the practice trials (i.e., clockwise response would be correct). To ensure participants properly learned the task, they only proceeded to the main experiment block after achieving 100% accuracy on the practice block for pairs separated by relatively large and suprathreshold distances (i.e., C vs. L6, C vs. L8, and L2 vs. L8). Feedback on accuracy was provided during practice blocks; in the main experiment participants were informed about the number of missed trials (including trials in which they reacted too slowly) at the end of each block.

### B. Results

Sigmoid functions were adequate fits to data reference points C, L1, R1, L2, R2, L3, R3, R4, L5, and L5 for all 6 participants. At other reference points, one or more participants performed at or below chance level (1 participant at reference points L4, R6, and R7, 2 at L6 and L7, 3 at R8, and 4 at L8). As it is impossible to estimate the 75% threshold for a flat line, those data were removed from the analysis. MAA values are illustrated in FIG 3 and summarized in TABLE I. The average MAAs across participants are in the range of 0.95° to 2.38° at reference points within about 32° off the midline (C, L1-L4, and R1-R4; see TABLE I). MAAs increased at more lateral locations. On the left, average MAAs were 4.46° for both L5 (50.41°off the midline) and L6 (69.09°off the midline), and were 2.81° and 8.48° for homologous right-sided locations (R5 and R6 respectively). These values are all consistent with published values^15,16^. We tested even more lateral locations, L7/R7 and L8/R8, located at 79.55°and 90.00° off the midline respectively. Performance at these locations was highly variable. At L7/R7, thresholds could be calculated for all 6 participants, and were 5.84°/7.94°. At L8/R8, values were based on data from two (L8) or three (R8) participants, and one of the participants included for L8 was an outlier with a threshold of 36.24 degrees. Therefore, these lateral data are probably not reliable.

**FIG. 3.**
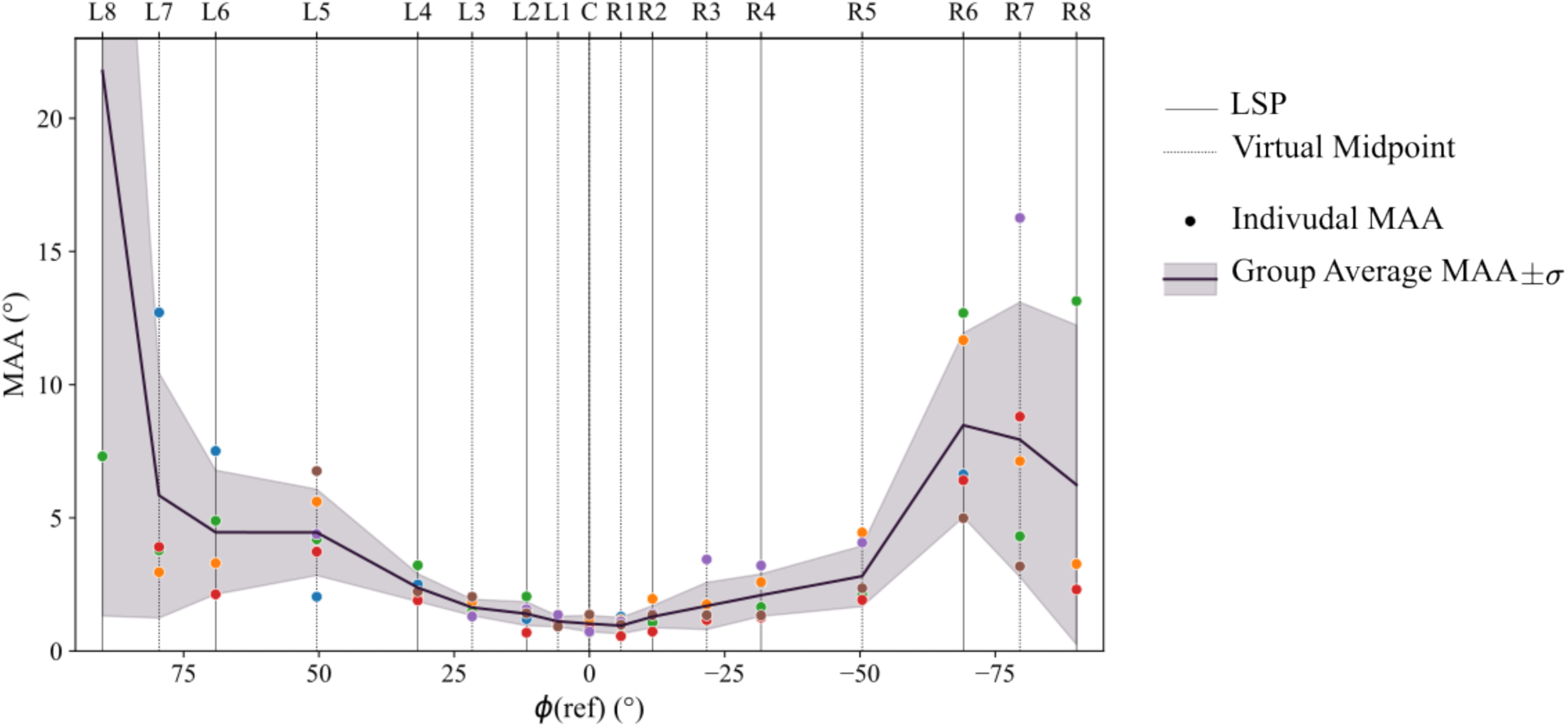
(color online) The horizontal MAAs for sound sources reproduced with 9^th^-order ambisonics. The line shows average values across participants; individual 75% thresholds are shown with dots color coded for each participant. The shaded region illustrates one standard deviation. (One individual’s MAA at −90° (L8) was 36.24°; this value is not shown in the figure.)

In the rotated conditions, when participants were facing reference points L5 and R5, the average MAAs were 1.20°±0.08° (N=6) at L5 and 1.17°±0.12° (N=6) at R5. (FIG 4.B). These values do not differ from the frontal MAA obtained at C (three-condition repeated-measures ANOVA; F(2,10) = 0.8275, p = 0.4650). As the LSP density at L5 and R5 is much less compared to the density at reference point C and humans are best in localizing sound sources in front of them, the similarity of frontal MAAs at all these locations indicates the homogeneity of ambisonic reproduction precision across the surface space, regardless of the density of the LSP array at that location.

**FIG. 4.**
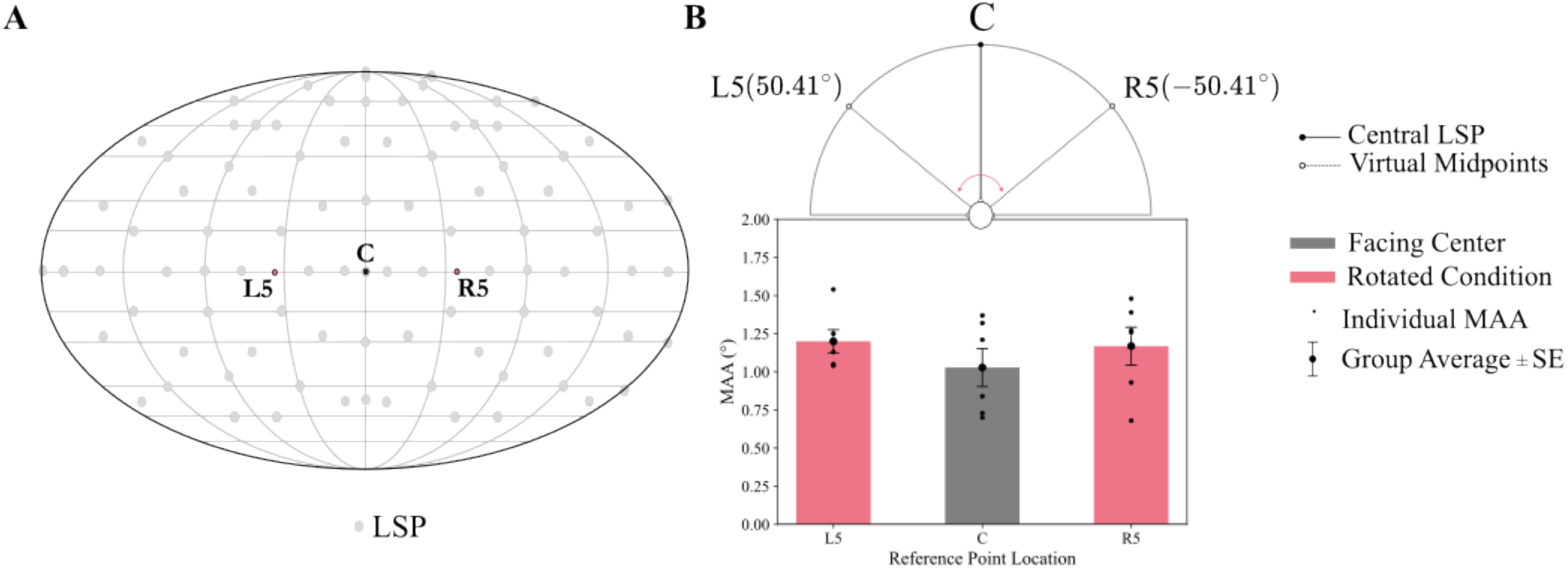
(color online) A. A flattened illustration of the LSP locations (light grey dots) in the array with reference points C, L5, and R5 (small dark dots). Note that C coincides with an LSP and is surrounded by other LSPs nearby (as close as ±11.64° horizontally), while L5 and R5 are ±18.69° azimuth and ±11.64° elevation away from the nearest LSPs. B. The midline MAA estimated at different locations with different LSP densities. Individual data are illustrated with small dots and the error bars show standard error of the mean.

In the post-experiment questionnaire, all participants reported perceiving noise bursts presented at lateral locations L6-L8 and R6-R8 as above or below the horizontal plane, and this was particularly true on the right side. Some participants used this illusory or artifactual elevation percept as an indication of location in the horizontal plane. For example, one participant had concluded that in trials at the R8 reference point, the noise burst that is perceived as higher was located clockwise relative to the other one. (This in fact was wrong and resulted in a flipped psychometric function.) The possible cause of the elevation percept is investigated further in the following experiments.

### C. Discussion

In this experiment horizontal MAAs were estimated for 17 reference points on the horizontal plane for wide-band noise bursts reproduced with 9^th^-order ambisonics. The estimated MAAs were consistently low around the midline and increased at more lateral locations. These results align well with previous reports of human spatial acuity tested in free-field settings: MAAs within the range of ±25° azimuth match with the values reported by Mills measured for 500 Hz and 1000 Hz pure tones^15^. At other locations, the estimated MAAs do not differ substantially from the Mills values except for L8 (90° on the left). Also, all frontal MAAs reported here (in both the original and rotated conditions) were smaller than those obtained in free-field testing with pink noise high-pass filtered at 4600 Hz, as reported by Strybel and Fujimoto^17^, possibly because of the greater sound information and low-frequency ITD cues available with broad-band noise. Our results demonstrate that sound sources reproduced with 9^th^-order ambisonic panning are sufficiently focal such that two adjacent sources can be resolved at or below the level of normal human spatial acuity at least for broadband sounds on the horizontal plane, within 50.41° off the midline. The MAAs obtained at lateral azimuthal locations (greater than, 50.41° from the midline) diverge from those recorded using free field methods^15,16^. The divergence of these may be due to the small number of samples, since several participants were excluded at these lateral locations. Another explanation for results in L7, R7, L8, and R8 would be the insufficient range of the test points; as we expected a lower acuity at these locations^15,23^, we tested locations further apart around these reference points (±12° for L8 and R8, and ±9° for L7 and R7 instead of the ±6° range for the other reference points). The main reason for data exclusion at these locations was the sigmoid model’s failure in estimating the data which was due to listeners’ inability to discriminate sound locations even at the most extreme test points. This suggests that a larger range of locations should be tested to provide a sufficient difference for location discrimination. We leave this as a caveat of our design as it was discovered after we analyzed the data.

Frontal MAAs were tested on the midline, both when the individual was facing reference point C, where the LSP density was highest, and also when the individual was facing L5 and R5 (see FIG 4), regions with the lowest LSP concentration. Obtained thresholds did not differ, implying that the simulation precision is homogeneous across regions with different LSP densities. This observation reassuringly indicates that sound sources are reconstructed with a spatial focality and sharpness that is truly location independent – at least with the 9^th-^order ambisonic panning used here. This confirms theoretical predictions about the advantage of higher-order ambisonics over VBAP^6^.

Finally, we investigated the anecdotal reports of elevation perception, which were unexpected. We first made sure that the simulation codes did not send any commands to reproduce sounds sources at any elevation level other than 0°. Then, three authors (NZ, BM, and IJ) and another person (DQ) sat in the AudioDome and listened to a block of the task at L8. We agreed that the virtual sound sources were sometimes heard from different elevation levels. This illusion disappeared when we rotated to face L8 in the middle of the block. This observation motivated Experiment 2, in which we quantified the localization cues for virtual sound sources rendered with higher-order ambisonics.

## III. EXPERIMENT 2

In Experiment 1, participants’ performance dropped substantially at lateral sound sources, more than expected based on previous free-field measurements^15,16^. For example, at *±*90° (reference points L8 and R8), data for at least half of the participants were at chance, or below chance, at all test points. Participants in Experiment 1 also reported hearing sounds from lateral locations as above or below the horizontal plane. These observations suggest that the spectral profile of the sounds was distorted and was providing artifactual cues to elevation. Given that elevation perception relies on high-frequency monoaural spectral information^1,24^, we suspected that the ambisonics algorithm or implementation (erroneously) resulted in the creation of notch filtering in the higher part of the spectrum.

To investigate whether elevation perception at lateral locations was perhaps a result of artifactual localization cues, we first evaluated spectral and ITD and ILD information for ambisonic sound sources as recorded through a Head and Torso Simulator (HATS) placed in the same position and facing the same direction as the participants in Experiment 1 (FIG. 5). Then we estimated the spectrum, ITD, and ILD values for wideband-noise sources presented from horizontal LSP locations using SC presentation, VBAP, and ambisonic panning. Then, the values were estimated for the low-frequency and the high-frequency portions of the spectrum and were compared across the two bands. As SC only applies the LSP frequency response to the audio files it serves as the “ground truth” for measurements. Hence, to identify errors in the spectral content of reproduced sound localization cues, ITD, and ILD values derived from recordings of sound presented using VBAP and ambisonic panning were compared to those using SC presentation.

**FIG. 5.**
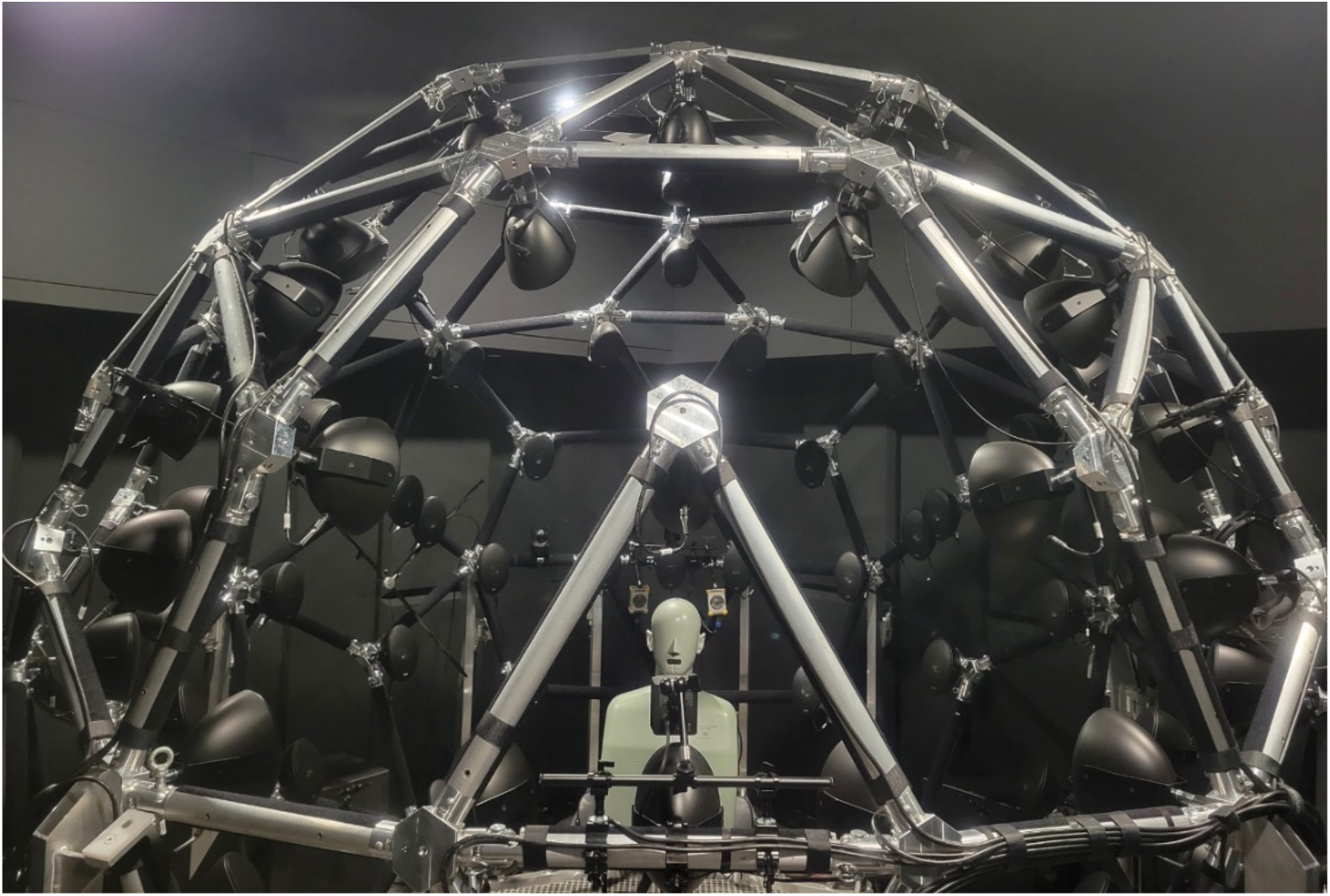
(color online) The HATS manikin in the measurement position in the AudioDome.

### A. Materials and Methods

The HATS type 4128C produced by Brüel & Kjær (Denmark) was used in this experiment. This manikin simulates the acoustic properties of an average adult’s upper body, head, and ears. The HATS was placed on a tripod at the position of a listener in the AudioDome and faced towards the origin of the system with the ears at 0° elevation level aligned with the ±90° azimuth line (FIG.5).

A sinusoidal sweep (chirp signal) was presented from target locations while the acoustic content was recorded by two half-inch microphones located in the HATS built-in ears; to estimate binaural and monaural cues as they are heard by an actual (average) human. The chirp signal was a broad-band linear frequency sweep starting from 0 Hz to 22050 Hz sampled at 44100 Hz rate with a duration of 22.05 s (1000 Hz/s) and a constant amplitude of 78.1 dB SPL (Equation 2).

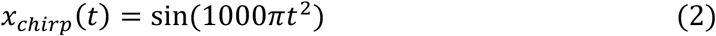

The sweep was presented from all the reference points tested in Experiment 1 (TABLE I) with VBAP and ambisonic panning modalities, and SC presentation where possible (where source location coincided with an LSP; i.e., reference points C, L2, R2, L4, R4, L6, R6, L8, and R8). For each method, sweeps were presented 4 times at each location, starting from R8 in counterclockwise order to L8. The sweeps were interleaved with 2 s periods of silence, to ensure that any reverberations from the previous sweep presentation had ceased, and that reproduction commands and triggers were transmitted properly. Data for all three reproduction methods were recorded in one session in the following order: SC presentation, VBAP, and ambisonic panning. The HATS position and orientation did not change during the recording.

Because the audible frequency range is known to be from 20 Hz-20 kHz^2^ and our system’s bandwidth is limited to 90-20730 Hz, we restricted the frequency range of all the following analysis to 100 Hz-20 kHz.

#### 1. Spectral content estimation

The power spectrum of the chirp signal, and of each recording, was estimated using the *pspectrum* function in MATLAB^25,26^ with a frequency resolution of 200 Hz and leakage of 0.85. To estimate the HATS transfer function for each presentation/panning method, at each target location, in each ear, the spectra of corresponding recordings were divided by the chirp signal’s spectrum and averaged across the 4 repetitions. To measure the difference between the original and recorded spectra for high and low frequencies, the spectrum was divided into two bands: a low-frequency band from 100 Hz to 4000 Hz, and a high-frequency band from 4000 Hz to 20000 Hz. The 4000 Hz boundary was chosen based on the equation introduced by Ward and Abhayapala^6,13^; this equation^1^ derives the minimum ambisonic order required to accurately reproduce a planewave with a particular wavelength within a sphere of a particular radius at the center of the array. Using this equation, our 9^th^-order system is capable of accurately reproducing frequencies up to 4000 Hz within a sphere of ∼12 cm radius at the center. (Male heads average 56.7 cm in circumference^27^, which, if approximated to a circle yields a radius of 9 cm. Thus 12 cm is a conservative estimate that also allows for some error in head position.) Then, the HRTF amplitude spectra of sounds recorded from VBAP and ambisonic panning were compared against SC presentation HRTFs using the gain-difference score ΔG as described in Equation 3^28^. In Eq. 3, |HRTF_panning_(f)| represents the HRTF magnitude of the ‘panning’ method at frequency ‘f’, and ‘f_L_’ and ‘f_H_’ represent the lower and the higher bound of the estimation frequency band, respectively.

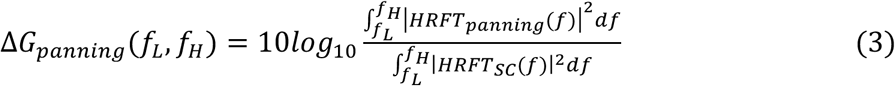

#### 2. ITD estimation

ITDs are more dominant for frequencies below 1 kHz^1,3,4^ and thus any reconstruction error at higher frequencies are not important. However, in the frequency range of 1000-1500 Hz neither ITD nor ILD cues dominate localization^1,3,4^, therefore to assess the general quality of ITD reconstruction, the 1000-1500 Hz range with some margin was included in the lower frequency portion of the spectrum for ITD estimations. For ITD measurement, the spectrum was again divided into two portions: but this time the lower frequency band extended from 100 Hz to 1800 Hz, and the higher frequency band from 1800 Hz to 20000 Hz. To estimate the ITD within each range, the portion of the recording that corresponded to each frequency range (based on the chirp signal presentation timing) was taken. Then this portion of the signal was temporally upsampled with an FFT padding factor of 10 to increase the temporal resolution of the cross-correlation function used in the next stage. Finally, the ITD was estimated as the lag amount that maximized the cross-correlation between the left and right channel recordings (delayed in the right channel relative to the left). ITDs were reported as the average of the 4 values estimated for each location with each reproduction method.

#### 3. ILD estimation

To estimate ILDs, the Fourier transform of the signal in both ears was estimated with the *fft* function in MATLAB^25,26^. As in the spectral content analysis, a boundary of 4000 Hz was used to split the spectrum range of 100 Hz-20 kHz into low-frequency and high-frequency bands. The sum of squared values of the Fourier coefficients of frequencies within each band indicated the spectral power of that band. The ratio of the spectral power in the left channel over the spectral power in the right channel was used to estimate ILDs in dB. ILDs were estimated for each measurement separately and the average across the 4 repetitions were used to calculate an ILD value for each reference point and reproduction method.

### B. Results

The HRTF for the left and right channels of the HATS for sound sources from horizontal LSP locations is shown in FIG 6.A for SC presentation and ambisonic panning methods. This panel, in combination with the gain-difference scores, ΔG, for the two frequency bands illustrated in FIG 6.B, shows that the ΔG is larger at higher frequencies than at lower frequencies, and, overall, ambisonically rendered sounds have lower power than SC sounds. (VBAP is theoretically equivalent of SC presentation where the sound sources coincide with an LSP location^8^, therefore VBAP HRTF magnitudes are not included. In panel B the ΔG between VBAP and SC is shown and approximates 0 everywhere, confirming the equivalence of VBAP and SC presentation methods for veridical LSP locations.)

**FIG. 6.**
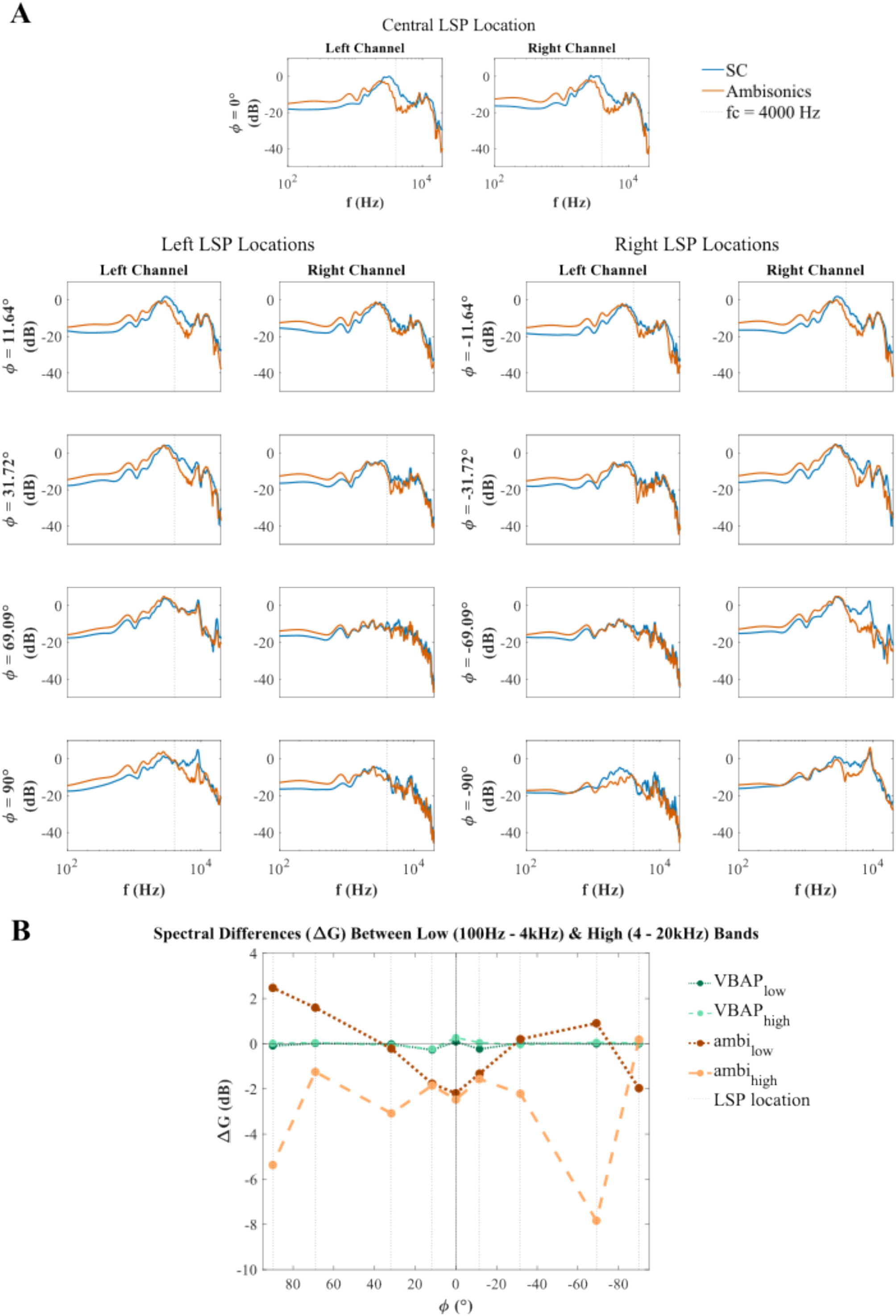
(color online) A. The HRTF magnitudes (left, and right) for sounds reproduced at LSP locations on the horizontal plane in the AudioDome. Only SC presentation and ambisonic panning methods are shown. The top row shows the spectra of sound sources presented on the midline (reference point C). On the other rows, the left pair of panels shows the spectra of response to presentations at L2, L4, L6 and L8, in the left and right ears. The right pair of panels show the spectra of response to presentations at R2, R4, R6, and R8. B. The gain-difference score (ΔG), which compares the magnitude spectra of SC and other presentation methods, is shown for the low frequency (100-4000 Hz) and high frequency (4000-20000 Hz) regions in the channel ipsilateral to the source location (for φ = 0° the average of both channels is illustrated). F_c_ = 4000 Hz was set as the boundary frequency that splits the spectrum. ΔG for VBAP - SC illustrates the noise ceiling of the score - the fact that its values are around 0 dB confirms the equivalence of VBAP and SC for these reference point locations.

For all the reference points tested in Experiment 1, the average ITD and ILD values were estimated for the three sound reproduction methods (only LSP locations for SC) as illustrated in FIG 7. A When the sound source was located on the horizontal midline (φ = 0°), ITD values were exactly 0 μs for all reproduction methods, and all repetitions. This confirms that the HATS was positioned properly in the center of the array during the experiment. The average low-frequency (100-1800 Hz) ITDs for SC and VBAP followed a monotonic trend from reference point L8 (–90°) through the midline (C) to R8 (+90°), and ranged between ±703 μs. ITD values were symmetrical between left and right. ITDs measured from VBAP presentations were equal to those of SC ITDs at LSP locations as theoretically expected. For ambisonics, the average low-frequency ITDs ranged from −658 μs to +681 μs with a monotonic trend from reference point L8 (–90°) through the midline (C) to R8 (+90°). ITDs were slightly different between the left and right hemifields (maximum difference ∼ 23 μs) at L3/R3, L6/R6, L7/R7, and L8/R8 pairs. The average low-frequency ITDs for VBAP and ambisonics were equal for L7/L8 and R7/R8 pairs. Average high-frequency (1800-20000 Hz) ITDs for SC also followed a monotonic trend within the range of ±726 μs, and had equal magnitudes on the right and left except at the L4/R4 pair where they differed by ∼23 μs. Average high-frequency ITDs for VBAP again matched the values of SC at LSP locations (except for at R8 where, they differed by 5 μs) and were symmetrical on the left and right, except for the L8/R8 pair which differed by 5 μs. At locations between LSPs, high-frequency ITDs derived from VBAP data diverged from the monotonic trend and were not always symmetrically equal in magnitude. ITDs measured in the high-frequency region for ambisonic presentation ranged from - 840 μs to749 μs, with a non-monotonic trend at the front (L2 to R2); the magnitudes of these ITDs were symmetrically equal up ±51°, and the trend diverged from SC ITD values on the sides (L6-L8 and R6-R8). The standard deviation (across four recordings per reproduction method, per reference point) of all measured ITD values was 0 μs, except for the high frequency band VBAP ITD at φ = −90° which had a standard deviation of 11.34 μs.

**FIG. 7.**
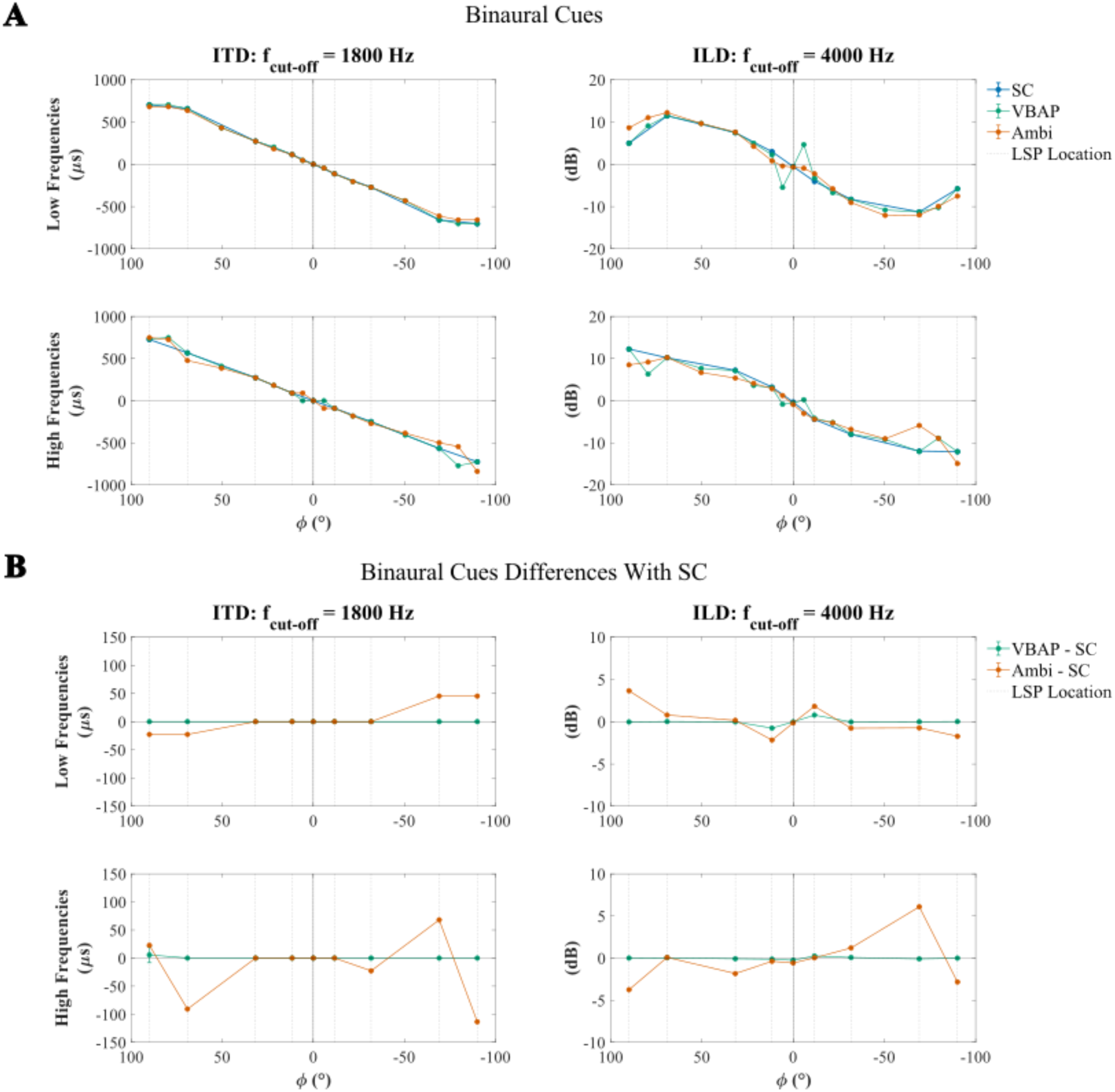
(color online) A. The average ITD (left column) and ILD (right column) values in low (top row) and high (bottom row) frequency bands across different reproduction methods for sound sources at reference points across the horizontal plane. The boundary frequencies to split low and high bands were 1800 Hz for ITD and 4000 Hz for ILD panels. Data for locations between LSPs are available only for VBAP and ambisonic panning. B. The average difference between estimated ITDs and ILDs for VBAP compared to SC and for ambisonic panning compared to SC at LSP locations (for better visualization of the small differences). In both panels error bars represent standard errors.

Average low-frequency (100-4000 Hz) ILDs for SC were within the range of ±11.5 dB with a monotonic trend within ±69.09° azimuth angle and a lower magnitude of ±5.8 dB at ±90°. The average low-frequency ILD was −0.55 dB when the source was at the center (reference point C). (Note that ITD was 0 μs at this point indicating that HATS was probably not poorly placed, so this small low-frequency ILD may result from different shapes of the left and right ears of the HATS.) At symmetrical locations, these values were relatively equal in magnitude (less than 1.1 dB different). For VBAP, the average low-frequency ILDs closely matched (less than 0.8 dB different) with SC at LSP locations but diverged from the SC trend at locations between LSPs, particularly at L1 and R1. Average low-frequency ILDs for VBAP were symmetrical on the left and right (less than 1.8 dB different). For ambisonic sources, average low-frequency ILDs were close to SC across the horizontal plane (less than 2.2 dB different) except at L8, where they differed by 3.7 dB. Average high-frequency (4000-20000 Hz) ILDs for SC-presented sounds ranged from −12.10 dB to 12.25 dB, and were bilaterally symmetrical (less than 1.8 dB different), and trended monotonically from reference point L8 (–90°) through the midline (C) to R8 (+90°). High-frequency ILDs for VBAP matched SC values (less than 0.3 dB different), and were relatively symmetrical (less than 2.6 dB difference between left and right homologous locations). VBAP high-frequency ILD measurements diverged from interpolated SC data between LSPs, particularly at L1, R1, L7, and R7. Finally, high-frequency ILDs measured from ambisonic presentation were close to those measured for SC presentation between 69.09° and −50.41° azimuth locations (less than 1.9 dB different) and had symmetrical magnitudes (less than 1.8 dB difference between left and right within ±50.41° range), but diverged significantly and asymmetrically from SC data (and interpolations) at more lateral locations. Finally, the standard deviation of the four ILD estimates did not exceed 0.07 dB at any location for any reproduction method.

Because the ITD and ILD differences between the SC and other modalities are small relative to the scale of the actual values, they are shown in panel B of FIG. 7 to provide a better visualization.

### C. Discussion

Frequency responses to broadband sweeps were measured from the left and right ear canals of a head and torso simulator (HATS), positioned in the AudioDome the same way as participants in Experiment 1, for all reference points tested in that experiment. SC presentation at the frontal LSP location (φ = 0°) produced responses in the left and right channels that are similar to each other and to frontal HRTF magnitudes previously reported^1^. At all locations, within each channel, the magnitude spectra of HRTF differed between ambisonic panning and SC presentation. Although the power of ambisonically rendered sound sources was generally less than that of SC sources, the difference in magnitude between the two reproduction methods was not homogenous across the spectrum. In the high frequency range (4000-20000 Hz), the power of ambisonically rendered sound was 1.5-8 dB below SC values for LSP-coincident reference points (the only exception was φ = −90°, where high frequency power matched between reproduction methods). In the low-frequency region (100 – 4000 Hz) the power difference between the two methods was in the ±2.5 dB range, which was overall lower than the power difference in the high-frequency region. Furthermore, the relative power of reproduction methods in the low-frequency band varied with the sound source location. (FIG 6.B)

Low-frequency (100-1800 Hz) ITDs for sources presented over SC followed a monotonic trend, from reference point L8 (–90°) through the midline (C) to R8 (+90°), within a range similar to previous reports; for example, Feddersen et al. demonstrated ITDs between ±670 μs over this range^1,29^. VBAP low-frequency ITDs matched these values at locations coincident with LSPs, as expected, but diverged from interpolated SC values at positions between LSPs. Low-frequency ITDs measured from ambisonically presented sounds were not only close to those of SC presented sounds, but they aligned better than did VBAP ITDs with the interpolated SC values at midpoint locations. For the higher frequency band (1800-20000 Hz), SC ITDs changed monotonically over the reference points from L8 to R8 as expected. VBAP and ambisonic ITDs diverged from interpolated SC values at locations between LSPs, and this divergence was more marked, and values were more asymmetrical between left and right, than for the low-frequency range. Fortunately, these higher-frequency ITDs are not relevant for perception and so these high-frequency ITD reproduction errors may not be perceptually salient. Thus our results indicate that neither VBAP nor ambisonic panning produce appreciable ITD errors (FIG 7, left column) and the ITDs accord well with those in the literature.

In the low-frequency region (100-4000 Hz), ILDs for SC-presented sounds are somewhat smaller than expected from published work. Feddersen et al^1,29^ demonstrated ILDs between 0-20 dB for frequency subbands below 4000 Hz, with lower frequencies having smaller ILDs. Our low values might be due to averaging across the band. VBAP values matched those for SC presented sounds at LSP locations but diverged from the interpolated values at locations between LSPs. In contrast, although low-frequency ILDs for ambisonically presented sweeps were somewhat different from SC values at LSP locations, they were overall closer to the interpolated SC data across all locations. In the higher frequency range (4000-20000 Hz) ILD values measured from VBAP presentation were similar to those for SC presented sounds at LSP locations but still diverged from interpolated SC values between LSPs. High-frequency ILDs for ambisonically presented sweeps matched SC values at frontal locations (C, L1-4; R1-4), but diverged significantly from the SC trend at the most lateral locations (L7-8, R7-8). High frequency ILDs measured from ambisonically rendered sweeps differed at homologous locations on the left and right. Given that ILD information used to localize sounds is derived from high-frequency components of sounds these differences between ambisonic and SC ILDs are likely to mislead human perceptual localization, especially when low-frequency ITDs are not present to down-weight high-frequency ILDs^4,30^(FIG 7, the right column). In general, binaural localization information is largely preserved for frequencies below 4000 Hz in the AudioDome, consistent with theoretical predictions^6,13^. ILD and HRTF cues at frequencies above 4000 Hz (such as monaural spectral cues used for elevation perception) may not be correctly reproduced.

Finally, the spectral content, ITDs, and ILDs for sound sources presented ambisonically at the most lateral locations (φ = ±69.09° and φ = ±90°) differ on the right and left sides (for example, look at the spectral power peaks at ∼8 kHz in FIG 6.A). These unexpected asymmetrical reconstruction errors are not observed when SC or VBAP reconstruction methods are used. The asymmetries are most visible in the magnitude spectra. To ensure that this was not an artefact of the recording setup or analysis method, the HATS was horizontally rotated 180° to face the back of the AudioDome and the chirp signals were presented at φ = ±90° locations with all three presentation methods again. The asymmetries in the estimated magnitude spectra were reversed in the rotated compared to the unrotated condition, indicating that the asymmetry results from the implementation of ambisonic panning in the AudioDome. (See supplementary FIG.S. 1.) This implies that there may be some asymmetry in the LSP or other hardware setup in the AudioDome, resulting in somewhat different spectral errors at very lateral locations – beyond φ = 69.09° - on the left and right sides at frequencies over 4000 Hz. Interestingly, other studies that simulated or tested ambisonic systems have also observed asymmetric reconstruction errors; Neal and Zahorik^12^ simulated SC, VBAP, and ambisonic reconstruction with different orders and estimated reconstruction error and the potential impact on psychoacoustic localization cues (total level, ILD, and ITD). Although not thoroughly discussed, they also report cue reconstruction errors that differ between the left and right side in their simulations, especially for higher-order ambisonics. Other examples can be found in figures 9.11 and 9.13 in the sound-field review chapter of “Immersive Sound” by Nicol^6^ that illustrate ITD and ILD estimates across different locations on the horizontal plane for a sound source at φ = 0° rendered using 19^th^-order ambisonic system. Again, asymmetry of reconstructed cues is observable, although not discussed and interpreted in the text. Ambisonic panning might be very sensitive to implementation details such as LSP calibration, adjustment, etc. Although it seems that reconstruction error asymmetry has not attracted much attention in the field, it could be an issue when ambisonic technology is used for psychoacoustic experimentation; erroneous conclusions about left-right asymmetry in human performance might be drawn when in fact the source of the asymmetry is, at least in part, in the implementation of ambisonics.

## IV. EXPERIMENT 3

During Experiment 1, participants anecdotally reported perceiving ambisonically presented sound sources, particularly those at lateral locations, as elevated. This was surprising, since all sounds were presented at locations in the horizontal plane. In Experiment 2, we determined that, at lateral locations, spectral information above 4kHz was distorted when sounds were rendered using ambisonic panning. Given that the spectral content in the 4-16 kHz range is important for sound source elevation perception^24^, these spectral differences might serve as unintended elevation cues. In Experiment 3, we aim to provide empirical evidence for the anecdotal reports in Experiment 1 and we hypothesize that they could be explained by the findings in Experiment 2. To achieve this goal, we had an independent group of listeners compare the perceived elevation of one of a pair of noise bursts, both presented at the same location: one reproduced with ambisonics and one presented through a single channel. We tested the same reference points as in Experiment 1, and participants were blind to the fact that sound sources were programmed to be collocated. Our hypothesis about systematic errors in the reproduction of high-frequency information creating artefactual elevation cues is investigated by comparing the degree to which lowpass, ambisonic sounds (i.e., without high-frequency cues) are heard as elevated above their SC controls, compared to wideband, ambisonic sources with high-frequency information, and thus artefactual elevation cues.

### A. Materials and Methods

Twelve (5 female) young (aged 22-30 years), healthy, normally hearing (thresholds measured audiometrically at both ears at octaves of 125 Hz up to 8 kHz; participants were excluded if their threshold exceeded 25 dB HL at any tested frequency in either ear), right-handed listeners with no history of neurological disorders completed the task. (Four listeners were excluded for the following reasons: two reported concussion history, one did not pass the audiometry test, and one reported during the debrief that they did not follow the eye fixation instructions.) Listeners had no experience with AudioDome experiments. The experiment setup and participant positioning were similar to Experiment 1 (except that no head tracking sensors were applied).

Participants completed an elevation discrimination task for reference points C, L4, R4, L6, R6, L8, and R8 as listed in TABLE I. On each trial, participants were presented with two noise bursts and were instructed to report the relative elevation of the second burst compared to the first one. The trials were presented in two conditions: in the ‘lowpass’ condition both noise bursts contained only frequency components below 4000 Hz (the boundary value for accurate acoustic environment reproduction with ambisonic panning as explained in Experiment 2^6,13^), and in the ‘wideband’ condition two wide-band noise bursts, with components between 0 and 22050 Hz (presented via the 90-20730 Hz bandwidth system), were presented. In both conditions the noise bursts were presented from the same location: one was presented with SC, and the other with 9^th^-order ambisonic panning. (Ambi_low_ vs. SC_low_, Ambi_wide_ vs. SC_wide_ trials). Twenty trials of each type were included for each reference point, with the SC-presented burst first on half the trials. Another forty trials per reference point were tested in which both noise bursts were presented with SC (SC_low_ vs. SC_low_, SC_wide_ vs. SC_wide_ trials) as filler trials (they were not analyzed later). To ensure participants stayed engaged, an additional 20 trials with large veridical elevation differences, both bursts presented using SC, were included at each reference point. The azimuth angle of sources in these trials was optimized to match the azimuth angle of the reference points, therefore, based on the AudioDome’s layout, the veridical elevation angle differences ranged between 61° and 94°. Thus, 100 trials were tested per reference point; 700 trials total.

As the task was identical to Experiment 1, except that the listeners had to judge elevation difference instead of horizontal differences, we used the same trial design to stay consistent with that experiment. The stimuli were generated using MATLAB^25^ with the same parameters as Experiment 1 (noise bursts duration, inter-stimulus pause duration, and the envelope were identical to Experiment 1). The lowpass noise bursts were originally generated as wideband (0-22050 Hz) and then, before the attack-decay envelope was applied, were filtered with an FIR lowpass filter with a passband frequency of 4000 Hz, a stopband of 4900 Hz, pass gain of 1 dB, stop gain of −80 dB, and a density factor of 20, designed and generated with MATLAB’s Signal Processing Toolbox^25,26^. Overall, 200 different noise bursts were generated and pseudo-randomly assigned to trials and intervals (we generated the noise burst audio sample sequence 200 times to enable our conclusions to generalize to other sounds that match the properties of ours. The same noise burst was assigned to both intervals of a single trial). The 700 trials were divided into five experimental blocks, each with an equal number of trials of all trial types at all reference points. The order of trial presentation was fully randomized. For each participant, stimulus assignment, trial order, and presentation method order within Ambi vs. SC test trials were pseudorandomized such that all noise bursts were presented an equal number of times at all reference points.

As in Experiment 1, lights were extinguished, and a red LED was placed at the front, on the horizontal plane at ear level for gaze fixation. Participants were instructed to report the relative elevation of the second noise burst compared to the first one (above/below) via keypress on a small keypad. At the beginning of the experiment, participants practiced the task on 14 trials in which both bursts were presented single channel and there was a large veridical elevation displacement between them (2 practice trials per reference point). They proceeded to the main trials once they passed a 75% accuracy threshold in the practice block (the block was repeated after clarification of the instructions if they performed with a lower accuracy). Feedback about accuracy was provided to participants only during the practice block. During the main experiment participants were informed about the number of missed trials (including trials in which they were too slow) at the end of each block keep them focused on the task. The data were collected within one session of 1 hour duration.

In the preprocessing stage, trials with no responses were removed from the data. Then, the proportion of trials on which the ambisonically presented sound was perceived as elevated was estimated for each participant, for all trial types, at each reference point.

### B. Results

We predicted that ambisonically rendered sounds with frequency components over 4 kHz would be heard as above or below the horizontal plane ompared to the same sound presented over a single channel on Ambi_wide_ trials, whereas ambisonically rendered sounds without such components would not (i.e., Ambi_low_ trials). In fact, in most cases the ambisonically presented sound was perceived to be below the SC sound. The average proportion of trials on which the ambisonically presented sound was perceived as elevated, as a function of reference point for both lowpass and wideband noise bursts is illustrated in FIG 8.A (Ambi_low_ and Ambi _wide_). These values are reliably below the 50% value expected by chance at C (t_low_(7) = −16.17, p < 10^-6^ & t_wide_(7) = −2.44, p < 0.05), L4 (t_low_(7) = −20.11, p < 10^-6^ & t_wide_(7) = −5.23, p < 0.05), R4 (t_low_(7) = −6.45, p < 10^-3^ & t_wide_(7) = −15.56, p < 10^-6^), and L8 (t_low_(7) = −4.87, p < 0.01 & t_wide_(7) = −3.23, p < 0.05). At R6 (t(7) = −2.51, p < 0.05) and R8 (t(7) = −6.07, p < 10^-3^) only the Ambi_low_ proportion is below the chance level. All other values do not differ from the 50% chance level (p > 0.05). It is possible that the timbral and power difference between ambisonically rendered and SC rendered sounds was erroneously interpreted as an elevation difference or the erroneous spectral cues generated a lower-than-horizontal percept, as participants reported the burst with less power (i.e., the ambisonically rendered burst) as lower in elevation.

**FIG. 8.**
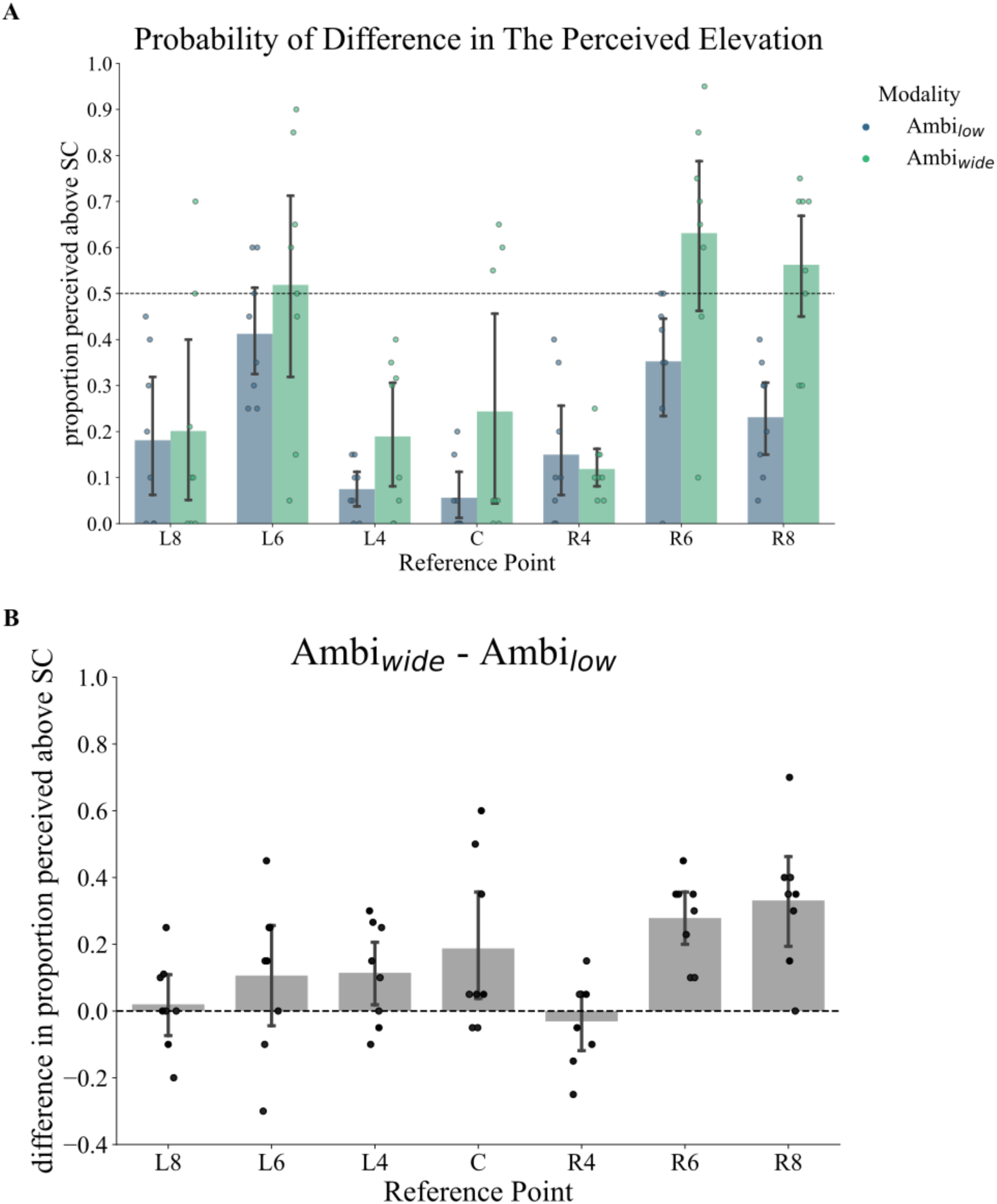
(color online) A. The average proportion of ambisonically rendered sounds perceived as elevated, compared to the SC reference interval for each of the two experimental conditions, Ambi_wide_ and Ambi_low_ B. The degree to which ambisonically rendered wideband noise bursts are reported as elevated (relative to their SC reference) compared to ambisonically rendered lowpass noise bursts (i.e., every set of two columns in panel A were subjected to the following arithmetic operation: Ambi_wide_ – Ambi_low_). Error bars in both panels represent the 95% confidence interval of the grand mean. Dots show individual participant data. Except at R4 which the grand mean is negative, all values are greater than zero, although only C, L4, R6 and R8 are significantly greater than zero, demonstrating a reliably greater tendency for wideband ambisonically rendered sounds to be perceived as elevated, compared to lowpass ambisonically rendered sounds, at these locations. The lack of significance at the other locations could be due to a lack of power.

Even though subjects exhibited a response bias towards reporting the SC presented sound as higher, we could still ask whether there is a difference in the degree to which wideband ambisonically rendered sounds are heard as elevated, relative to the lowpass ambisonically rendered sounds (which presumably contain fewer misleading localization cues). We therefore tested the difference between Ambi_wide_ –and Ambi_low_. This value was computed individually for each person and is shown in FIG 8.B. This value will be positive if Ambi_wide_ trials yield a greater proportion of elevation responses than do Ambi_low_ trials, as predicted. These values were indeed significantly positive at C (t(7) = 2.06, p < 0.05), L4 (t(7) = 2.12, p < 0.05), R6 (t(7) = 6.25, p < 10^-3^), and R8 (t(7) = 4.61, p < 10^-2^) reference points, but not at R4, L6, and L8.

### C. Discussion

In Experiment 3, we compared the perceived elevation of lowpass and wideband sounds, ambisonically reproduced at horizontal reference points, relative to the same sounds presented from a single LSP at the same location. Rather surprisingly, the ambisonically rendered sounds appeared to be generally perceived as below their SC controls. There is a perceptible timbral difference between the ambisonic and SC rendered sources, and the ambisonic sources also have lower power (as documented in Experiment 2). Participants may have interpreted these differences in the ambisonic sounds to indicate a source below the horizontal plane.

Despite this general response bias, we observed that a greater proportion of ambisonic sounds were perceived as elevated relative to the SC reference sound on wideband trials, compared to lowpass trials, at almost all reference points (FIG 8.A). This consistent effect suggests that the high-frequency components included in the wideband, but not lowpass, sounds were distorted by ambisonic presentation, contributing to perceived elevation as predicted. However, there was also more variability in response to wideband than lowpass trials. This is not surprising because each participant has their own HRTF, so individuals will interpret the spurious spectral cues present in the high-frequency components differently. Also, as illustrated in FIG 8.B, the effect of high-frequency components in the perceived elevation of ambisonic sources seems to be somewhat variable across different locations, being more pronounced at lateral locations on the right side. This is consistent with our observations in Experiment 2 of larger reproduction errors on the right side, compared to the left, possibly due to subtle asymmetries in the location or calibration of LSPs.

## V. CONCLUSIONS AND DISCUSSION

In this study, we assessed the fidelity of a 9^th^-order ambisonic LSP array for the reproduction of focal sound sources on the horizontal plane. In Experiment 1, we quantified the limits of human spatial acuity in the frontal half of the horizontal plane with wideband sound sources presented with ambisonic panning. Values for the minimal audible angle (MAA), the just noticeable difference in horizontal displacement ranged from 1.03 - 1.2°, for the horizontal midline location, similar to previous findings^15^. In fact, MAAs were similar to the existing literature for locations within the range of ±51° degrees azimuth, confirming that ambisonic panning in the AudioDome results in sources that are so focal that any blurring is below the threshold for human auditory spatial acuity. The recordings through a head-and-torso simulator in Experiment 2 demonstrated that ITD cues, estimated for ambisonic sources on the horizontal plane at frequencies up to 1800 Hz, are not distorted.

Our system is theoretically restricted to accurately reproduce plane-waves up to 4 kHz for a sphere of radius of 12 cm^6,13^, and indeed we noticed distortions in the frequency region above 4 kHz, which altered the spectral profile in ways that might affect localization. Spectral magnitudes in the range of ILDs were also affected, with left-right asymmetries that could create artefactual ILDs. In Experiment 3, we showed that ambisonic reconstruction errors in the frequency region 4000-20000 Hz are associated with a greater tendency to perceive the rendered sources as elevated, compared to similar sounds without components in this frequency region. Although for both behavioral experiments (1 and 3) we had a small number of listeners, we could capture highly robust results for conditions wherein we expected a consensus across participants: the frontal MAAs were highly precise and consistent and performance in the control condition in the elevation discrimination task was at a chance level with high statistical power. On the other hand, we observed high variability in lateral MAAs and elevation discrimination between SC and ambisonic sound sources which are most probably caused by the individual variability in interpreting the inaccurate localization cues by listeners. Such individual variabilities are due to the listeners’ HRTF variability. However, the small number of participants deserves to be acknowledged as a limitation of these experiments that could affect the conclusions and implications. Nevertheless we convincingly showed that the AudioDome can reproduce focal sources with a spatial fidelity that challenges the limits of human spatial acuity, particularly if high frequencies (above 4000 Hz) are omitted.

Based on these observations it seems best to limit the reproduced spectral content in the AudioDome to components below 4000 Hz for accurate spatial reproduction. For naturalistic studies of speech listening in the presence of competing talkers and other sounds, inclusion of information above 4000 Hz is desirable and so SC reproduction may be preferable.

Additionally, we showed that ambisonic panning reproduces sources more homogenously and reliably than does VBAP, at locations between LSPs. We also observe a general difference of reproduction error of high frequency components of sound between left and right symmetrical locations (here, high frequency is above 1800 Hz for ITDs and above 4000 Hz for ILD and spectral cues); although this asymmetry might be only specific to our system, there is evidence of this phenomenon in other studies simulated ambisonic rendering^6,12^. While its basis might be related to the sensitivity of the ambisonic implementation to LSP array calibration or positioning, we could not find any evidence for asymmetrical misalignment of LSPs or asymmetrical errors in encoding and decoding the spatial audio. Therefore, the causes of asymmetrical reproduction errors are unknown and future work is required to reveal the causes of this observation.

This project is the first study that aimed to utilize the AudioDome to test human perception, and it is also a pioneer in conducting psychophysical experiments using a higher-order ambisonics system, demonstrating that sources rendered with a 9^th^ order ambisonic system are sufficiently focal such that any blurring is below the threshold of human auditory spatial resolution. Although there are issues with ambisonic rendering of components above 4 kHz, this research lays the groundwork for future experiments on naturalistic human auditory perception.

## SUPPLEMENTARY MATERIAL

See supplementary material at [URL] for the comparison between HATS magnitude spectra to chirps from lateral LSPs in both ears (FIG.S. 1) and Experiment 3 participants’ performance at control trials (FIG.S. 2).

## ACKNOWLEDGMENTS

The authors would like to thank Dr. Derek Quinlan and Karsten Pravoslav Babin for their invaluable help and support with experiment hardware setup and programming. We thank the Canadian Foundation for Innovation (CFI) for funding the AudioDome, and the Natural Sciences and Engineering Research Council (NSERC) for research funding.

## AUTHOR DECLARATIONS

## Ethics Approval

The experiments presented in this manuscript were approved by The Western University Non-Medical Research Ethics Board. Informed consent was obtained from all participants.

## DATA AVAILABILITY

The data that support the findings of this study are available from the corresponding author upon reasonable request.

## Footnotes

1 *N* = ⌈*kr*⌉ with 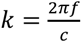 where *N* is the minimum ambisonics order required to reproduce a reliable plane wave with frequency of *f* in a sphere of radius *r* in an environment with sound speed of *c*.

