## Supplementary material for "Focality of sound source placement by higher (9^th^) order ambisonics and perceptual effects of spectral reproduction errors": Supp figures

At the end of Experiment 2, the HATS was rotated horizontally 180° so we could measure the effect of physical orientation of the HATS on the recording content; it is possible that the asymmetrical reproduction errors are due to asymmetries in the HATS structure (the model's pinnae for left and right are not perfectly symmetrical as natural pinnae). If this hypothesis is true, then the response of the left and right channels should not change as we rotate the manikin and record the same sounds from ±90° azimuth coordinates on the horizontal plane. However, as observed in FIG S.1, this is true for SC presentation, but for ambisonics, the response in the left channel transfers to the right ear and vice versa, showing that the recording is reflecting the true spectral content of the reproduced sounds at the recording location.

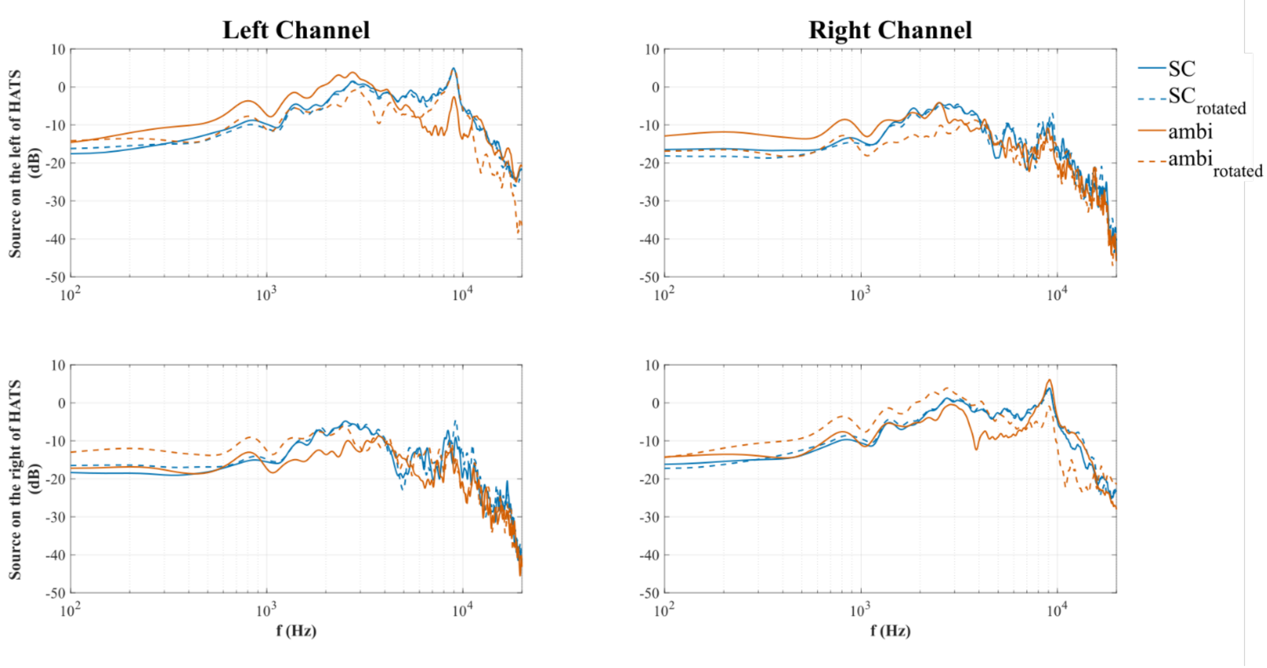


FIG.S. 1. (color online) The magnitude spectra of auditory responses to chirp signals presented from L8 and R8 (±90°) in both channels when the HATS is facing the center (φ = 0°) and when it is fully rotated to face the back of the AudioDome (φ = 180°).

In experiment 3, participants completed a set of control trials in which sound sources with the same frequency content were presented from the same location ($SC_{low}$ vs. $SC_{low}$ and $SC_{wide}$ vs. $SC_{wide}$). As there is no difference between the frequency content or location of these sounds and they are presented from a single LSP, participants' discrimination probability in these trials shows their bias. Ideally, the bias should be around 50% for this two-alternative choice that is within the confidence interval of average performance level in these control condition except for wide-band sounds at the location of R4. FIG.S. 2 illustrates this discrimination bias values that show low variability across participants.

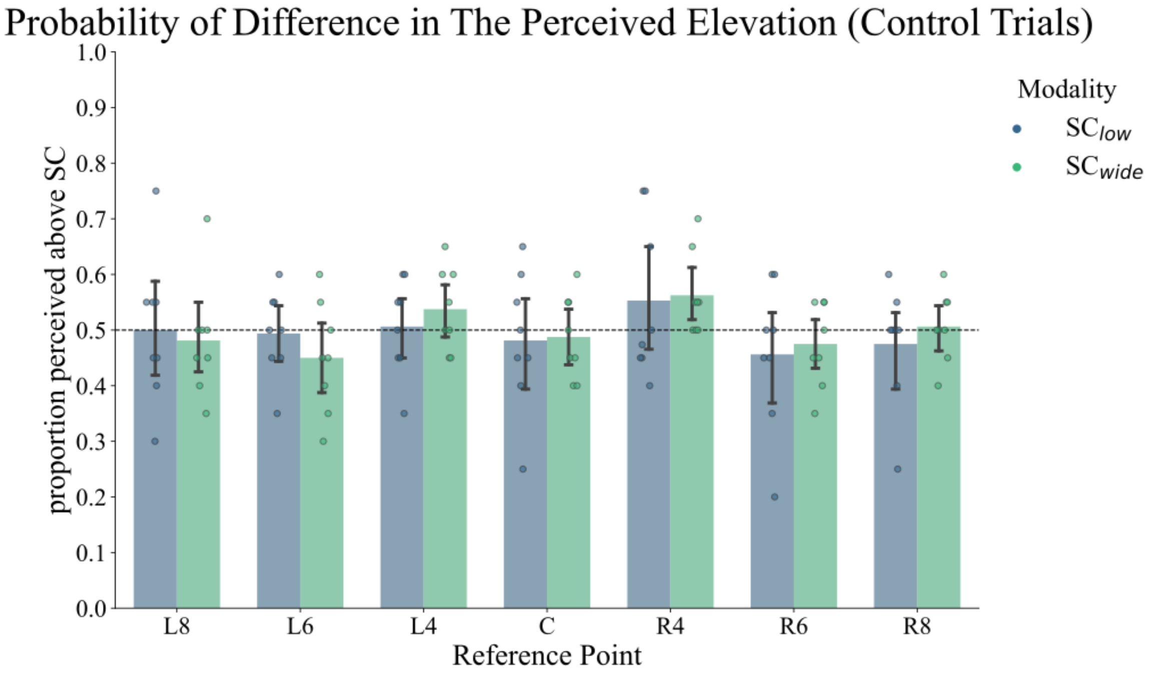


FIG.S. 2. The average probability of discriminating sound sources with both low and wide-band frequency content against the sources with the same characteristics from the same LSP on the horizontal plane, showing the listener's average bias in elevation discrimination.
